# Manganese-induced Parkinsonism in mice is reduced using a novel contaminated water sediment exposure model

**DOI:** 10.1101/541664

**Authors:** Dana M. Freeman, Rachel O’Neal, Qiang Zhang, Edward J. Bouwer, Zhibin Wang

**Affiliations:** Department of Environmental Health & Engineering, Johns Hopkins University, Baltimore, MD

## Abstract

The effects of heavy metals on human health have become an important area of study. For instance, acute manganese toxicity is known to induce Parkinsonism. Heavy metals including manganese enter the aquatic environment from both anthropogenic and natural processes. These metals accumulate within water sediments and their behavior is then dependent upon the sediment composition and phase. These metal-sediment interactions remain to be explored within *in-vivo* animal studies. To study the effect of these interactions, herein we successfully developed an exposure model in mice that encapsulates the aquatic microenvironment of heavy metals before exposure. Male and female *C57/BL6* mice were exposed to manganese contaminated sediment via their drinking water (Sed_Mn) or to manganese placed directly into their drinking water with no prior sediment interaction (Mn) for six weeks. Sediment interaction did not alter total manganese in drinking water (mg/L) or weekly manganese consumption (mg) in males (54.9±1.5 mg) or females (44.6±1.0 mg) over the six-week exposure period. We analyzed motor impairment, a common feature in Parkinson’s disease, using the beam traversal, cylinder, and accelerating rotarod behavioral tests. We observed Parkinson’s like deficits in motor control in both treatment groups as early as four weeks of exposure in males but not in females. Intriguingly, mice given water incubated with manganese spiked sediment (Sed_Mn) performed better overall compared to mice given manganese directly in water (Mn) despite having similar exposure in males and females. Male Sed_Mn mice compared to Mn mice had a 146% reduction in time to cross the beam traversal test (*p<0.05*), a 10% increase in rearing activity in the cylinder test (*p<0.05*), and a 14% increase in time remaining on the rotarod (*not significant*). Female Sed_Mn mice compared to Mn mice had no change in the time to cross the beam traversal test, a 36% increase in rearing activity in the cylinder test (*p<0.05*), and a 35% increase in time on the rotarod (*p<0.05*). Our study indicates that metal-sediment interactions may alter metal toxicity in mammals and introduces a new exposure model to test the toxicity of metal contaminants of drinking water.

## Introduction

Heavy metal contamination of drinking water poses a significant public health risk across the globe (EPA 2015). Exposure to heavy metals such as lead, arsenic, and manganese are associated with increased risks of various cancers, declined cognitive ability, and altered thyroid function (EPA 2015). The World Health Organization (WHO) has sought to mitigate health effects resulting from excess metal exposure via drinking water by setting global maximum contaminant levels (MCLs) (WHO 2011). Unfortunately, due to the challenges associated with climate change, aging infrastructures, and the lack of a comprehensive removal technique, many populations are exposed to levels of metals much higher than WHO guidelines (Chowdhury 2016).

Toxicology studies regularly use mouse models to gain mechanistic insights into heavy metal exposure and chronic illnesses. These studies model exposures by using invasive techniques such as oral gavage or by repeatedly dosing large concentrations of heavy metal salts directly into the water supply. A limitation of this method is the omission of complex metal-sediment interactions within water systems that may alter metal toxicity. Heavy metals accumulate within sediments and concentrations can exceed those in the water by three to five orders of magnitude (Bryan & Langston 1992). Metals can be transformed within these sediments into compounds with altered bioavailability and behavior (Nicolau 2006). While metal speciation and bioavailability in water is well understood; it remains challenging to study these reactions within complex microenvironments such as sediments (Bryan & Langston 1992, Islam 2015). The goal of this study was to develop an *in-vivo* exposure model that includes the effects of metal-sediment interactions to test the toxicity of heavy metals in water.

To our goal, we used manganese, a common water contaminant with well-studied toxic effects in animal studies. Manganese toxicity is associated with neurological dysfunction and Parkinson’s disease (Chartlet 2012, O’Neal & Zhang 2015, Bouabid 2016, Sarkar 2018). Excess manganese exposures in animals have been shown to disrupt mitochondrial function, induce neuroinflammation, obstruct neurotransmission, and damage the basal ganglia of the midbrain (Bouabid 2016, Sarkar 2018). Extensive neuronal death and tissue damage in the midbrain impairs motor function control and can be reliably measured in animals using various motor behavioral tests (Brooks & Dunnett 2009). Despite a robust animal phenotype, there is a lack of human data on the risks of Parkinson’s disease from the ingestion of manganese from drinking water (Chartlet 2012). Therefore, further understanding of the mechanisms occurring in the brain following manganese ingestion is needed. A recent study in plants observed reduced toxicity of manganese in water with the presence of silicates in the soil by reducing uptake into the cytoplasm (Blamey 2018). We hypothesized a similar mechanism may occur in mammalian systems. We exposed wild-type *C57/BL6* mice to manganese contaminated drinking water for six weeks by either putting manganese directly into the water (see methods-Mn) or by first incubating water with manganese contaminated sediment (see methods-Sed_Mn). The effects of different treatments on Parkinsonism in mice were determined by measuring altered motor function in three behavioral tests: the beam traversal test, the cylinder test, and the rotarod test. The aims of this study were (1) to determine if mice could be exposed to metals via contaminated sediment (2) to determine if incubating water with contaminated sediment changes the behavioral phenotype over time (3) to determine if there were sex dependent effects. Herein we report our mice exposed to manganese contaminated sediment produced a Parkinsonian phenotype in males but not females. We observed altered manganese toxicity in both males and females exposed to manganese contaminated sediment despite having been exposed to the same amount of total manganese (mg) as mice given manganese directly in water (Mn). We observed a trend of increased sensitivity of males to manganese treatment in both manganese treated groups (Sed_Mn and Mn) in at least two behavioral tests.

## Methods

### Chemicals and Manganese Water Preparation

Ottawa Sand from Restek (Bellefont, PA) was purchased as the sediment. The sediment control contained 1 kg of sand per 2L of Nestle Pure Life water in an autoclaved glass bottle. The contaminated sediment was spiked with manganese chloride (Sigma, St. Louis, MO) at 1 g MnCl2·4H2O per 1 kg of sediment. The concentration of manganese in sediment was selected based on our previous work analyzing heavy metal contamination of Baltimore harbor sediments using inductively coupled plasma mass spectrometry (ICP-MS) (Graham 2009; Wadhawan 2013). Water and spiked sediment were incubated out of light at room temperature two weeks before treatment began. The water was filtered from the sediment every three days before being given to the mice. The pH of the filtered water was monitored during the study and remained between 6-7 which is suitable for groundwater. The manganese chloride water solution was prepared fresh every three days at a concentration of 0.5 g/L.

### Animals and Treatment

Forty wild-type male and female *C57/BL6* mice ages 8-10 weeks were purchased from Jackson Laboratory (Bar Harbor, ME). Mice were housed 5 mice per cage on a 14-10 hour light-dark cycle in a AAALAC accredited facility. Mice were given food and water *ad libitum* and allowed to acclimate for 10 days prior to treatment. All procedures involving animals were performed under the guidance of the National Research Council’s *Guide for the Care of Laboratory Animals* (NRC 2010) and approved by the Johns Hopkins University Animal Care and Use Committee. Mice were exposed via their drinking water for six weeks. They were given freshly prepared manganese contaminated water every three days. Food and water intake was recorded bi-weekly. Body weights were recorded weekly.

### Manganese Detection Assay and Estimated Daily Exposure

Water filtered from the manganese sediment reaction was collected at the beginning and the end of the study and stored at −20°C. The detection of manganese in water sediment samples was completed using the sodium periodate oxidation method with instructions and reagents provided by the manganese test kit model MN-5 purchased from HACH (Loveland, CO). A standard curve was generated using three freshly prepared manganese standards at 100, 500, and 1000 mg/L. Three technical replicates for each standard and sample were measured by UV Vis absorbance at 525 nm in a 96 well clear bottom plate. The estimated daily exposure of manganese contaminated sediment water was calculated using the average concentration of the two unknown samples and the weekly water intake rate of each cage divided by the average body weight (kg) of each cage.

### Behavioral Tests

#### Rotarod Test

The accelerating rotarod test was used to assess motor coordination on the first and last day of treatment. The automated accelerating rotarod from Harvard Apparatus (Cambridge, MA) was generously loaned to us by the Jiou Wang laboratory at Johns Hopkins Bloomberg School of Public Health. Each mouse was trained one day prior to testing for both timepoints and given as many trials as needed to successfully balance on the moving rotarod for three runs with one minute of rest between runs. On test days, the mice were given three trials on the rotarod accelerating linearly from 4 rpm to 40 rpm for a maximum of five minutes. The latency to fall (seconds) was recorded for each mouse.

#### Cylinder Test

The cylinder test was used to assess balance and exploratory rearing behavior every two weeks. Mice were placed in a clear glass cylinder (diameter = 10 cm) under minimal light at the end of their dark cycle for a total of two minutes. Rears were defined as the lifting of one or both forelimbs above shoulder level and contacting the wall of the glass cylinder (Cannon 2009). The mouse must bring all forelimbs back to the ground before another rear was counted. Rears were counted by the handler at the time of testing and by a blind observer using a video recording of each test.

#### Beam Traversal Test

The beam test was used to assess balance and motor coordination every week. Mice were placed on the far end of a wooden rod (diameter = 1.6 cm) approximately 60 cm from a dark cardboard box containing the home cage bedding. In the first week before exposure, mice were trained one day prior to testing for baseline abilities. Training consisted of acclimating the mouse to the box and giving the mouse as many trials as necessary until it was consistently moving forward across the beam into the box. The mouse would be given approximately 30 seconds to rest in the box between trials. On testing days, if a mouse turned around on the beam or fell off the beam then that trial would be discarded. The mouse would be returned to the box before another trial was started. The mouse was video recorded, and the time for the mouse to cross the beam across three successful trials was reported by a blind observer.

### Biostatistics

One-way analysis of variance (ANOVA) was used to analyze differences in behavior between the four mouse treatment groups within each time period for both males and females. Two-way ANOVA was conducted to analyze the effect of time and sediment incubation on behavioral phenotypes expressed as the fold change compared to the water or sediment water control. Post hoc analysis was performed with both one-way and two-way ANOVA using the Tukey Honest Significance Difference test. The Student’s t-test was used to compare sex dependent effects on behavioral phenotypes expressed as the fold change compared to the water or sediment water control. Post hoc Bonferroni correction was used for significance of the Student’s t-test. For all analyses, *p<0.05* was considered significant.

## Results

### Manganese exposure does not alter food/water intake or weight

Food and water intake was monitored during the 10-day acclimation period and bi-weekly during the six weeks of exposure. The amount of food and water consumed by male and female mice remain unchanged during the study regardless of treatment group (Figure 1). Intriguingly, the control female cage had a significantly higher water intake rate compared to the other female cages that began before treatment and continued throughout the study (Figure 1d). We do not believe reduced water intake in the other cages was due to manganese exposure. The weight of each mouse was recorded weekly. No mouse lost more than 20% of their body weight during the exposure period and the change in weight of each mouse was not different among treatment groups (Figure 2). Overall male mice gained an average of 0.8 g over the six-week period while female mice gained 1.47 g (*p≤0.01, Student’s t-test*).

**Figure 1.**
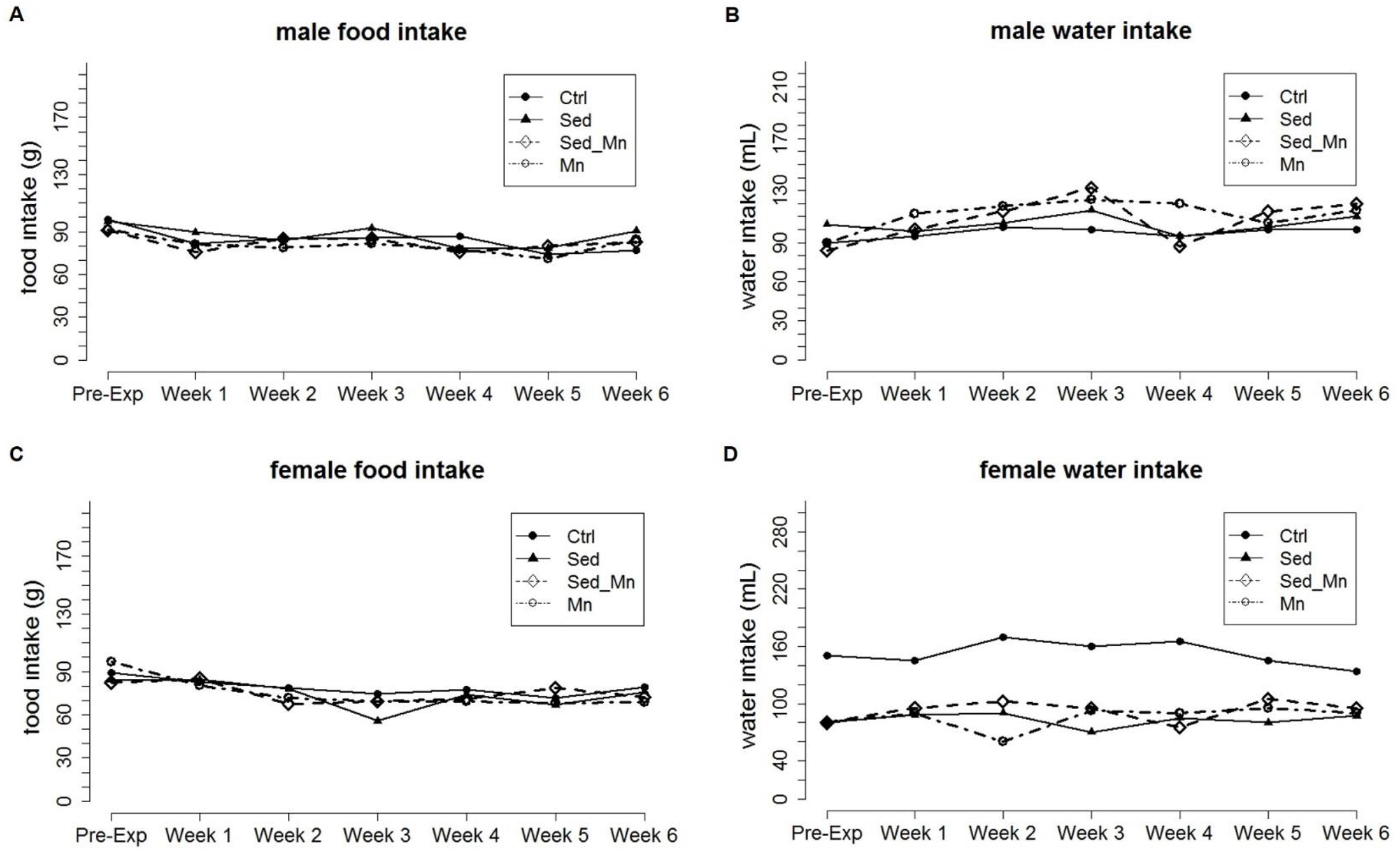
Weekly food and water intake changes in response to manganese exposure. A-B) Male weekly food and water intake rate by cage from week 0 (pre-exposure) to week 6. C-D) Female weekly food and water intake rate by cage from week 0 (pre-exposure) to week 6.

**Figure 2.**
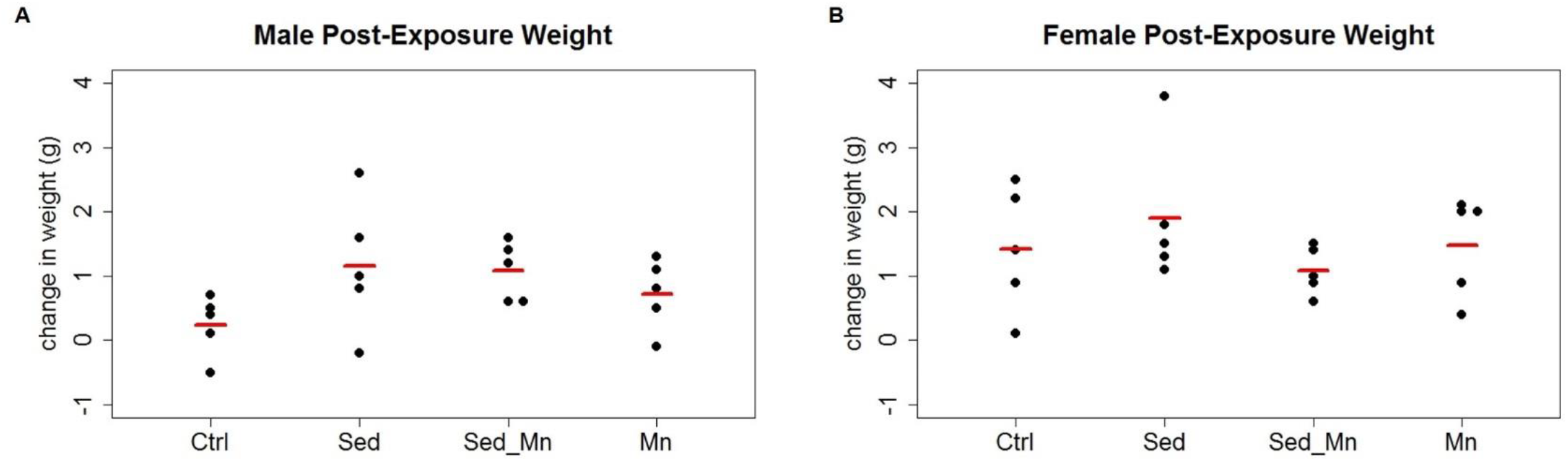
Mouse weight change after six weeks of manganese exposure. A-B) Male and female total change in weight at week 6 from week 0 (pre-exposure). Red bar indicates mean for each group (n=5). Significance >0.05 with ANOVA.

### Manganese sediment interaction does not alter total manganese in drinking water and males consume more total manganese than females

The effect of sediment incubation on manganese content was determined from filtered water collected at the beginning and end of the six-week exposure period. The abundance of manganese in the water was 446 and 473 mg/L, respectively. There was no significant difference in the total amount of manganese consumed by the manganese water (Mn) and the manganese sediment (Sed_Mn) cage for males or females (Figure 3a). However, female treatment groups (Mn and Sed_Mn) did consume less total manganese than males. We used the average body weight for each group to determine if the estimated daily exposure (mg/kg*day) was significantly different for males and females. Male Sed_Mn treated mice had a lower estimated daily exposure than female Sed_Mn treated mice (54 mg/kg*day and 62 mg/kg*day, respectively). Mice treated with manganese directly in the water received the same estimated daily dose of approximately 61 mg/kg*day regardless of sex (Figure 3b).

**Figure 3.**
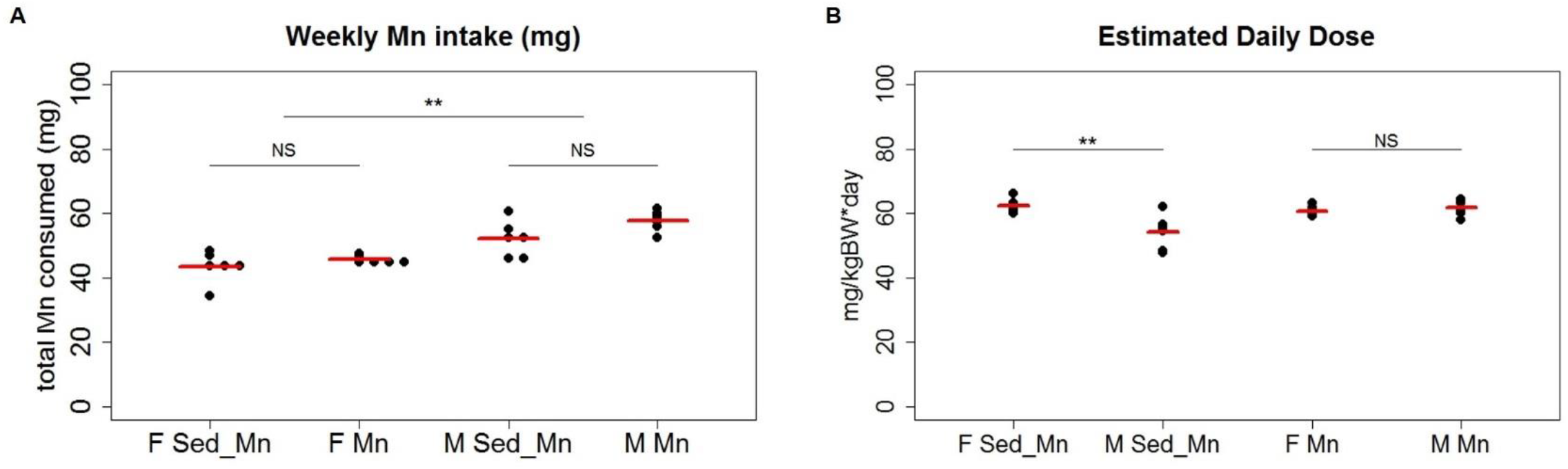
Weekly manganese intake and estimated daily dose did not change between treatment groups. A) Weekly manganese intake (mg) of female and male mice exposed to sediment water (Sed_Mn) versus manganese water (Mn). Red bar indicates mean (n=6 weeks). There was no difference in the total manganese consumed between Sed_Mn and Mn treatment groups (ANOVA, p>0.05, not significant NS). Male treatment groups consumed higher amounts of manganese than female groups (ANOVA, p<0.01, **). B) The estimated daily exposure dose for all groups. Males exposed to sediment water (Sed_Mn) had a lower daily exposure than females (ANOVA, p<0.01, **). Male and females exposed to manganese directly in water (Mn) both had an estimated daily exposure of 61 mg/kg.

### Sediment interaction reduces the effect of manganese on beam test performance in males

We examined the effect of sediment interaction on the development of Parkinsonism by using the beam traversal test to examine motor coordination and balance weekly during the six weeks of exposure. We recorded the average time over three trials for each mouse to cross the beam to the box containing the home cage bedding. No significant changes were detected in males or females until week six of exposure (Figure 4). At week six, only males exposed to manganese directly in water (Mn) took longer to cross the beam than the control mice (Figure 4a). The effect of both length of exposure and manganese sediment interaction on beam test performance was observed in males. Manganese sediment interaction significantly reduced the time to cross the beam in week four and six of exposure (Figure 4b) (*two-way ANOVA, p≤0.05*). The exposure time between week four and week six did not significantly increase the observed effect of sediment interaction. Neither female treatment group had reduced performance on the beam test during the study and there was no observed effect of sediment incubation on the average time to cross the beam. Therefore, our results indicate a possible gender-dependent behavioral phenotype in response to manganese.

**Figure 4.**
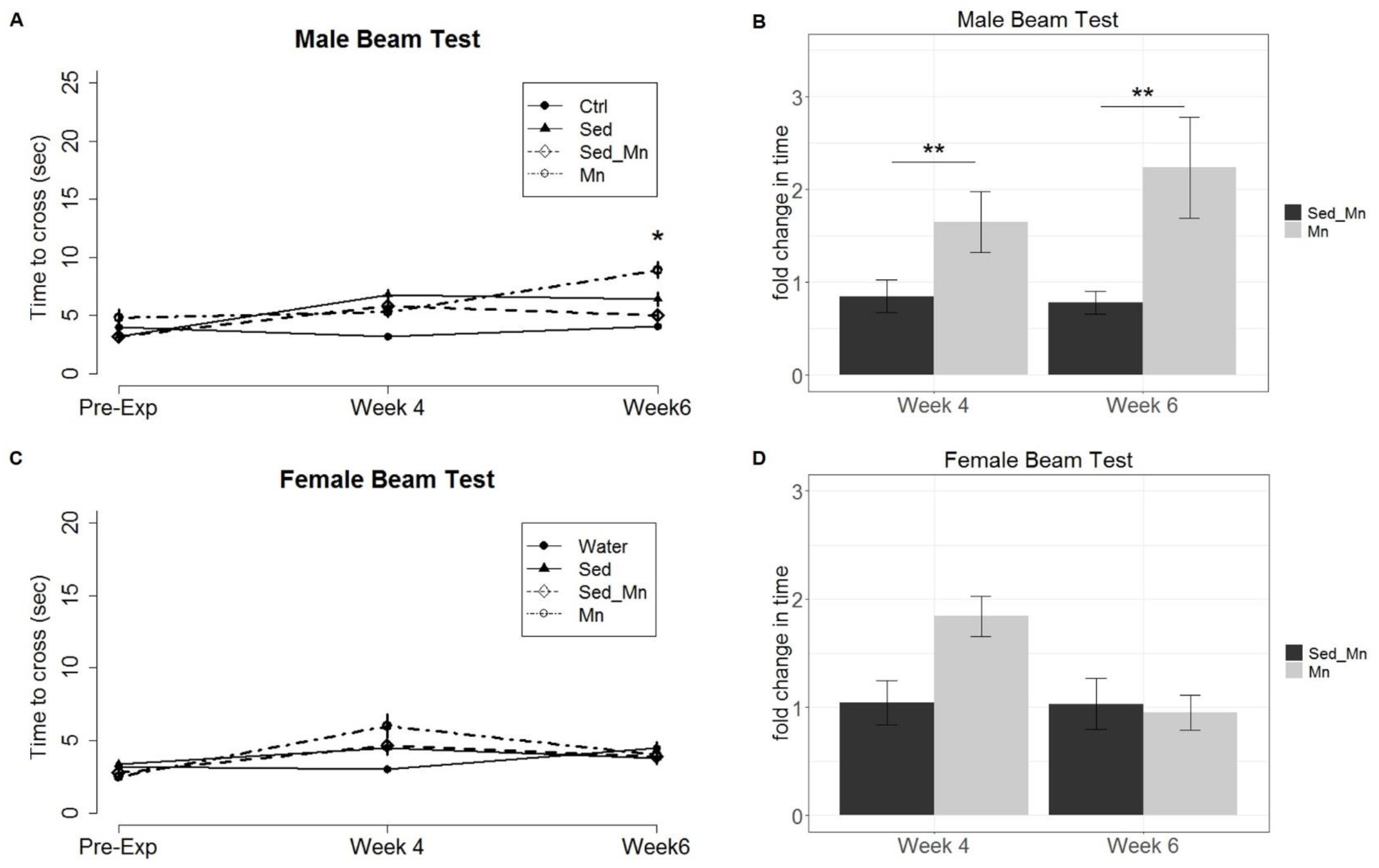
Examination of the average time to complete the beam traversal test between manganese treatment groups at week 4 and week 6 revealed that sediment interaction decreased effect on motor coordination in males. A) Average time to cross beam (seconds) ±SEM at week 0 (pre-exposure), week 4, and week 6. Male Mn but not Sed_Mn mice took significantly longer to cross the beam than control mice (ANOVA, p<0.05, *). B) Average fold change in time to cross beam ± SEM compared to respective controls (Mn/Ctrl; Sed_Mn/Sed). Fold change in time was altered with manganese sediment interaction but no significant effect of exposure duration was observed (two-way ANOVA, p<0.01, **). C) Average time to cross beam (seconds) ±SEM at week 0 (pre-exposure), week 4, and week 6. Female mice exposed to manganese did not take longer to cross the beam than control mice. D) Average fold change in time to cross beam ± SEM compared to respective controls (Mn/Ctrl; Sed_Mn/Sed). Fold change in time was not significantly altered in either treatment group and there was no observed effect of exposure duration.

### Sediment interaction reduces the effect of manganese on cylinder test performance in males and females

We examined rearing behavior every other week using the cylinder test. Rearing describes the lifting of the forelimbs onto the walls of the glass cylinder and allows us to analyze motor coordination using natural exploratory behavior without prior training. We observed a decrease in rearing behavior in male and female Mn groups at week four and six but not Sed_Mn (Figure 5a, 5c). The effect of both length of exposure and manganese sediment interaction on rearing behavior was observed. Manganese sediment interaction significantly reduced the number of rears counted in males and females at week four and six (Figure 5b, 5d) (*two-way ANOVA, p≤0.05*). The length of exposure had a significant effect on the rearing behavior in males indicating an increased effect of sediment interaction with time (*two-way ANOVA, p<0.01*). This was not observed in females.

**Figure 5.**
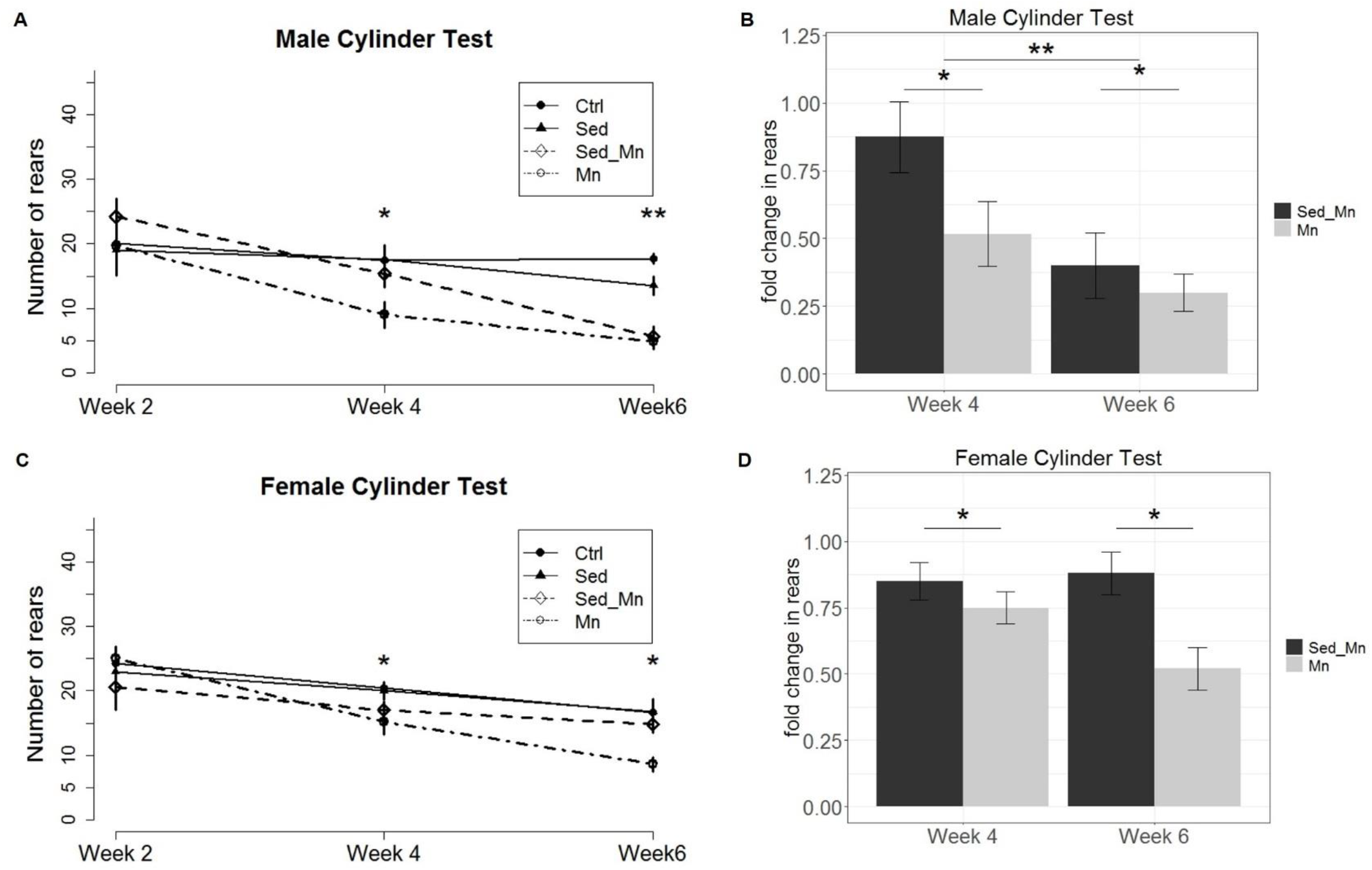
Examination of exploratory rearing behavior between manganese treatment groups at week 4 and week 6 revealed that sediment decreased effect on rearing activity in males and females. A) Average number of rears ±SEM at week 0 (pre-exposure), week 4, and week 6. Male Sed_Mn and Mn mice had significantly reduced rearing behavior compared to control mice at week 4 (ANOVA, p<0.05, *) and week 6 (ANOVA, p<0.01, **). B) Average fold change in rearing ± SEM compared to respective controls (Mn/Ctrl; Sed_Mn/Sed). Fold change in rearing was altered with manganese sediment interaction (two-way ANOVA, p<0.05) and with exposure duration (two-way ANOVA, p<0.01, **). C) Average number of rears ±SEM at week 0 (pre-exposure), week 4, and week 6. Female Mn mice had significantly reduced rearing behavior compared to control mice at week 4 (ANOVA, p<0.05, *) and week 6 (ANOVA, p<0.05, *). D) Average fold change in rearing ± SEM compared to respective controls (Mn/Ctrl; Sed_Mn/Sed). Fold change in rearing was altered with manganese sediment interaction (two-way ANOVA, p<0.05) but not with exposure duration.

### Sediment interaction reduces the effect of manganese on rotarod test performance in males and females

We examined balance and motor coordination at the end of the study using the accelerating rotarod test. This test measures the amount of time a mouse can balance on a rotating rod and is one of the most common tests used to quantify neurological deficits in rodents (Brooks & Dunnett 2009). We observed a decrease in time on the rotarod in both the male Mn and Sed_Mn groups when compared to the control mice *(ANOVA, p<0.05)* (Figure 6a). Manganese treatment did not significantly decrease the time on the rotarod in females (Figure 6c). Manganese sediment interaction reduced the fold change in time on the rotarod in both males and females but was only significant in females *(Student’s t-test, corrected p<0.05)* (Figure 6b, 6d).

**Figure 6.**
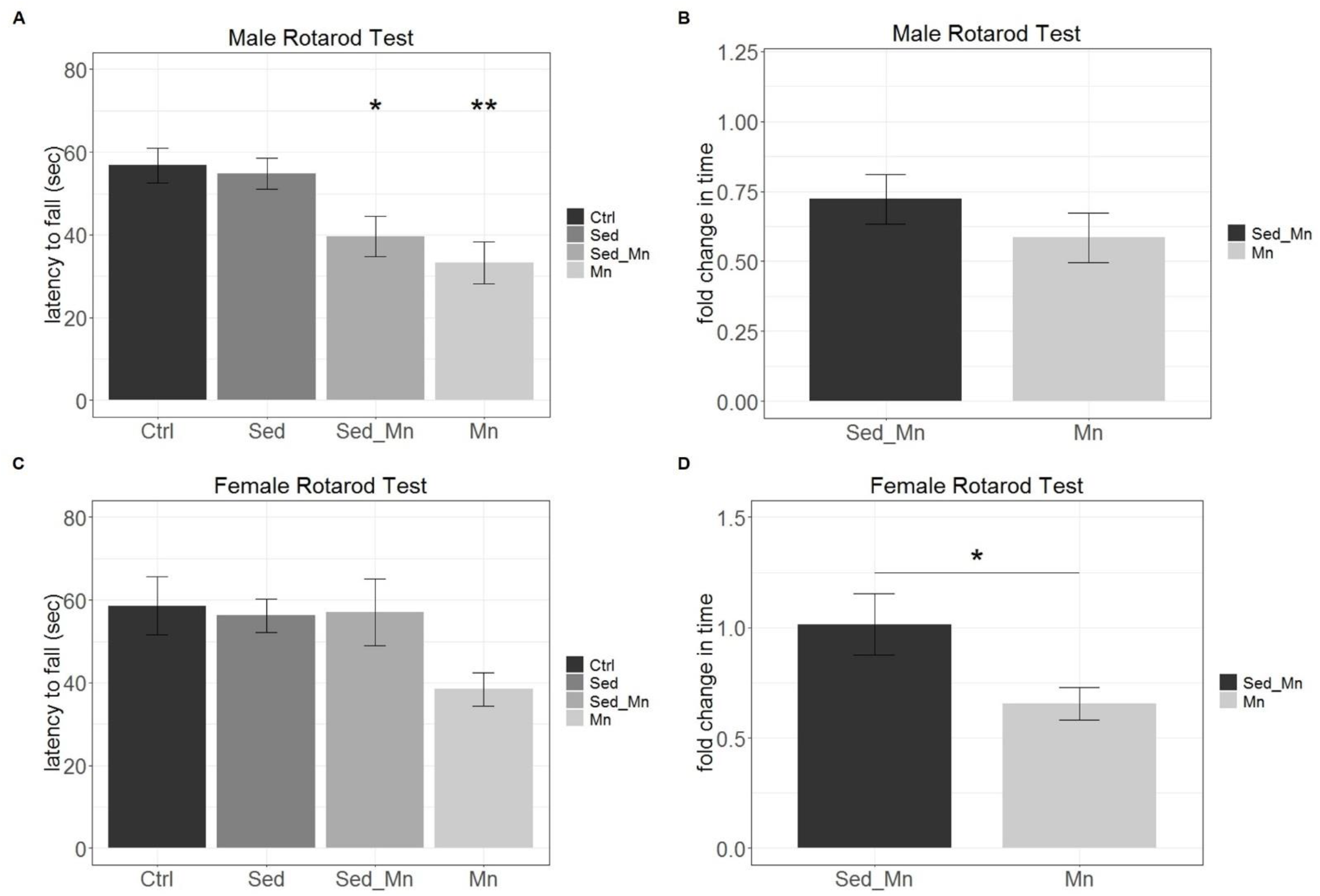
Examination of the average time on the accelerating rotarod between manganese treatment groups after six weeks of exposure reveals decreased effect on motor coordination in females. A) Average time to fall (s) ±SEM at week 6. Male Sed_Mn and Mn mice remained on the rotarod for significantly less time than control mice at week 6 (ANOVA, p<0.05, *, p<0.01, **) B) Average fold change in time ± SEM compared to respective controls (Mn/Ctrl; Sed_Mn/Sed). Fold change in time was altered with manganese sediment interaction (Student’s t-test, p<0.01). C) Average time to fall ±SEM at week 6. There was no significant difference in the average time to fall in manganese exposed female groups (ANOVA, p>0.05). D) Average fold change in time ± SEM compared to respective controls (Mn/Ctrl; Sed_Mn/Sed). Fold change in time was altered with manganese sediment interaction (Student’s t-test, p<0.05).

### Males performed worse than females in at least one of the three behavioral tests in both manganese treated groups

We observed differences in behavioral outcomes for males and females exposed to manganese during the study. We analyzed motor deficits at week six using the fold change in performance compared to the treatment group’s respective controls (Mn/Ctrl; Sed_Mn/Sed) (Figure 7). In the beam test, males took a significantly longer time to cross the beam than females in the Mn exposure group (*Student’s t-test, corrected p<0.05*). However, in the Sed_Mn exposure group, males and females performed the same and were not different than controls. In the cylinder test, males had reduced rearing behavior than females exposed to manganese in both groups but only the Sed_Mn group was significantly reduced from females after p-value correction (*Student’s t-test, corrected p<0.05*). In the rotarod test, males spent slightly less time on the rotarod than females in both exposure groups, but these were not found to be significant.

**Figure 7.**
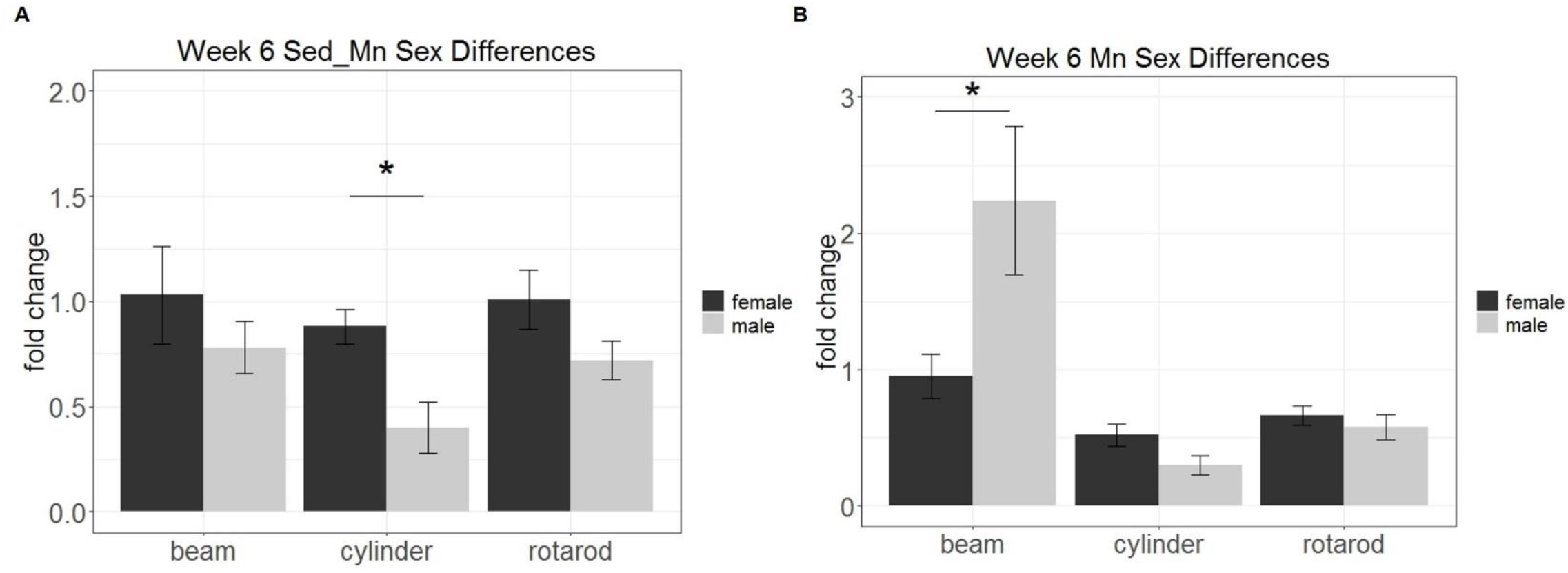
Examination of sex differences in behavioral testing reveals increased sensitivity of males in at least one behavioral test in both manganese treatment groups. A) Average fold change of the Sed_Mn treated male and female mice compared to the sediment control (Sed) ± SEM at week 6 (Student’s t-test, corrected p<0.05). B) Average fold change of the Mn treated male and female mice compared to the control ± SEM at week 6 (Student’s t-test, corrected p<0.05).

## Discussion

In this study we used manganese to test our proposed exposure model because of its well-studied mechanisms in the brain and its reproducible motor phenotype in mice. We chose a commercial sediment for this pilot test to avoid convoluting the end behavioral outcome. The commercial sediment was made predominately of silica which is a major constituent of watershed and marine sediments. Manganese is known to leach to minerals such as silicates in the watershed and mineral content in water sediments has been of interest to study increased risks of neurodegeneration (Choi 2006; Chartlet 2012). The sediment used in this model was sterile and had a consistent pH but potential changes in these factors on metal toxicity will still need to be studied in the future by using more complex sediments in the exposure model.

We selected an environmentally relevant sediment concentration of manganese of 1000 mg/kg sediment based on our previous work in the Baltimore City harbor (Graham 2009; Wadhawan 2013) and expected 0.5g/L as the maximum concentration of total manganese in the water after incubation. This concentration was also used for the Mn treatment group in which manganese water was prepared fresh every three days. We predicted that 0.5 g/L would be sufficient for this model based on another study that used the same dose in the drinking water of *C57/BL6* males and found significant deposition of manganese in the brain and a measurable motor phenotype (Krishna 2014). The concentration was not expected to be lethal given numerous studies using concentrations ≥ 1g/L to study neurotoxic effects in rodents (Chandra 1981; Anderson 2008; Avila 2008; Alsulimani 2015). We closely monitored food intake and weight during the study to verify that behavioral changes were due to neurotoxic mechanisms and not acute systemic failure. We observed no significant reductions in food intake or weight among mice in the study (Figure 1 and 2).

It was unclear whether water incubation with sediment would affect the taste of the water and alter the mouse rate of ingestion. We recorded weekly water intake and found that mouse drinking behavior was not significantly altered from the pre-exposure acclimation period suggesting that mice did not seem to experience any taste aversions (Figure 1). We used the sodium periodate oxidation method and UV absorbance to detect total manganese content in water after filtration of contaminated sediment. The average total manganese was 0.46 g/L indicating that manganese was not simply degraded or being filtered out of the drinking water. We used the manganese concentration and water intake to compare manganese ingestion between treatment groups. We found no differences in the amount of manganese consumed weekly within male or female manganese treated mice (Figure 3). To determine if males and females received a similar dose of manganese we calculated the estimated daily exposure using the weekly manganese ingested divided by number of mice per group (n=5) and the average weight per group. We did not observe any change in the estimated daily exposure between male and female Mn treatment but did observe a reduction in the daily exposure of Sed_Mn males compared to Sed_Mn females (Figure 3).

We successfully exposed mice to manganese via contaminated sediment and produced a Parkinsonian phenotype in males as early as four weeks of exposure. Male Sed_Mn mice performed significantly worse than male controls in the cylinder test and the rotarod test. They performed better than male mice treated with Mn directly in water in all three tests after six weeks of exposure despite having the same total amount of Mn in the drinking water. The female Sed_Mn group did not develop a phenotype after six weeks of exposure. However, female Mn mice only performed worse than controls in one test suggesting a possible resistance of females to manganese treatment. This is supported by the literature in which multiple studies have reported an increased sensitivity of males compared to females possibly due to estrogen inhibition of the NFKB inflammatory pathway involved in neurodegeneration (Moreno 2011; Gillies 2014).

We hypothesized that metal-sediment interactions may alter metal toxicity from ingestion of drinking water by changing metal behavior. Manganese bioavailability and uptake across physiological membranes depends heavily on its oxidation state. Manganese speciation can also alter metal behavior in intracellular fluids. Intracellular divalent cations including Mn2+, the most common free cytosolic manganese species, can interact with proteins and effect the function of metabolic enzymes. Divalent manganese may also complex with amyloid fibrils, prion proteins, and Lewy body inclusion bodies that are highly prevalent in neurodegenerative diseases including Parkinson’s disease (Choi 2006; Chartlet 2012). These complexes are associated with the observed perturbation of intracellular degradation systems such as the ubiquitin-proteasome complex but whether divalent metal ion complexes cause or are an effect of misregulated protein oligomerization remains uncertain.

Current animal studies analyzing heavy metal toxicity do not consider the microenvironment of the metals in the aquatic environment or the complex interactions that occur within sediments. Our study provides an *in-vivo* model to expose mice to water contaminants that simulates sediment interactions. We supported our hypothesis that sediment interactions alter metal toxicity in mammals. Future studies should include more complex sediments including those with microorganisms that can metabolize metals and affect metal behavior in water. Additionally, phenotypic tests may be altered and should be selected based on predicted metal contaminants and their target organ systems. Heavy metal contamination of drinking water is a global public health issue and mechanistic insights into the effects of metal contaminants is needed for the protection of human health. Our exposure model can be used to improve toxicity testing on single metal contaminants and metal mixtures that are present in water systems.

## Acknowledgements

This project is made possible with support from the EHE Chen Award (ZW and EB) and JHU Catalyst Award (ZW), as well as student Frank Bang Award (to DMF). Experimental disposals and salaries were partially supported by the U.S. National Institutes of Health (R01ES25761, U01ES026721 Opportunity Fund, and R21ES028351) to ZW. We also thank Dr. Jiou Wang and student Reham Aljumaah for their generous help with the accelerating rotarod.

## Data Availability

The data used to generate this report are available upon request from the corresponding author.

